# Endocytosis suppresses stochastic collapse in fibroblast-macrophage circuits under shared resource competition

**DOI:** 10.64898/2026.05.27.728330

**Authors:** Ken-ichi Inoue, Yusei Ishii, Masanori Hariyama

## Abstract

Interdependent multicellular circuits must maintain stable coexistence despite competition for shared environmental resources. Fibroblast-macrophage circuits represent a conserved signaling architecture in which fibroblasts produce colony-stimulating factor 1 (CSF) to support macrophages, whereas macrophages produce platelet-derived growth factor (PDGF) to support fibroblasts.

Previous analytical models proposed receptor-mediated endocytosis as a stabilizing negative-feedback mechanism, but these formulations assumed spatial homogeneity and independently assigned carrying capacities.

Here, we constructed a spatial agent-based fibroblast-macrophage circuit model using PhysiCell to investigate how PDGF and CSF endocytosis regulate circuit stability under explicit competition for shared oxygen and space. Fibroblasts and macrophages competed for common environmental resources supplied by spatially distributed capillary sources, allowing carrying capacity to emerge dynamically from local resource competition.

Across nine enhancer conditions spanning fourfold variation in PDGF and CSF signaling strength, heterotypic coexistence remained broadly achievable regardless of endocytic activity. In contrast, endocytosis strongly suppressed stochastic circuit failure. This stabilization depended critically on macrophage CSF uptake, whereas broad ranges of fibroblast PDGF uptake produced comparable outcomes, generating a permissive stabilization landscape along the PDGF uptake axis. Mechanistically, excessive CSF signaling drove macrophage overexpansion, depletion of shared resources, and eventual fibroblast extinction.

Importantly, despite fundamentally different carrying-capacity assumptions from previous analytical models, both frameworks converged on the same systems-level conclusion: stabilization of the macrophage-supporting CSF axis is substantially more critical than stabilization of the PDGF axis.

These results identify endocytosis as a robustness mechanism that suppresses catastrophic failure in interdependent multicellular circuits under shared-resource competition without requiring precise parameter tuning.

## Introduction

Multicellular tissues are maintained by interdependent signaling circuits in which distinct cell populations exchange trophic factors to coordinate survival, proliferation, and repair. Among these, fibroblast-macrophage interactions constitute a conserved multicellular circuit observed across multiple physiological and pathological contexts, including wound healing, fibrosis, tumor stroma, and tissue-resident macrophage niches. In this circuit, fibroblasts produce colony-stimulating factor 1 (CSF1) to support macrophages, whereas macrophages produce platelet-derived growth factor (PDGF) to support fibroblasts [1]. This circuit architecture is conserved across multiple mammalian tissues, with fibroblast-derived CSF1 supporting tissue-resident macrophages demonstrated independently in the spleen, intestine, and skin [2–4].

Analytical studies of fibroblast-macrophage circuits proposed that stable coexistence requires asymmetric negative feedback on the macrophage-supporting CSF axis [1]. In these formulations, fibroblasts were treated as near carrying capacity whereas macrophages remained proliferative, creating a “spring-and-ceiling” architecture in which stabilization depended critically on suppressing excessive CSF accumulation. Subsequent theoretical work demonstrated that receptor-mediated endocytosis can function as a robust stabilizing mechanism by reducing growth-factor accumulation through ligand uptake [5].

However, these analytical models assume spatial homogeneity and independently assigned carrying capacities. In living tissues, fibroblasts and macrophages coexist within heterogeneous microenvironments in which oxygen, nutrients, and physical space are shared resources rather than cell-type-specific constraints. Under such conditions, carrying capacity may emerge dynamically from local competition rather than being assigned independently to one cell population. Spatial competition can fundamentally alter multicellular dynamics and generate emergent behaviors absent from mean-field approximations [6].

In particular, interdependent signaling circuits may be vulnerable not only to deterministic instability, but also to stochastic collapse arising from local resource depletion and probabilistic extinction events. Whether endocytosis stabilizes fibroblast-macrophage circuits under these spatially explicit and stochastic conditions remains unclear.

To address this problem, we constructed a spatial agent-based fibroblast-macrophage circuit model using PhysiCell [7]. Unlike previous analytical formulations, fibroblasts and macrophages in our model competed explicitly for shared oxygen and space supplied by spatially distributed capillary sources. We systematically varied PDGF and CSF enhancer strengths together with fibroblast PDGF uptake (Puptake) and macrophage CSF uptake (Cuptake) across stochastic spatial terrains.

We found that endocytosis did not substantially enhance coexistence probability once heterotypic clusters formed. Instead, endocytosis selectively suppressed stochastic circuit failure. This stabilization depended strongly on macrophage CSF uptake, whereas broad ranges of fibroblast PDGF uptake produced comparable outcomes, generating a permissive stabilization landscape along the PDGF uptake axis. Importantly, despite fundamentally different carrying-capacity assumptions from previous analytical models, both frameworks converged on the same systems-level conclusion: stabilization of the macrophage-supporting CSF axis is substantially more critical than stabilization of the PDGF axis.

## Materials and Methods

### 1. Agent-based simulation framework

All simulations were performed using PhysiCell (version 1.13.1), an open-source physics-based agent simulation framework for multicellular systems biology [7]. Diffusion and uptake of extracellular substrates were solved using the BioFVM finite-volume diffusion framework [8].

Simulations were conducted in a two-dimensional 2,000 × 2,000 µm domain using a BioFVM mesh spacing of 20 µm. Diffusion time steps were 0.01 min, cell mechanics time steps were 0.1 min, and phenotype updates were performed every 6 min. Each simulation was run for 30,240 min (21 days).

The model was designed to investigate stochastic stability of interdependent fibroblast-macrophage signaling circuits under explicit competition for shared spatial resources.

Complete PhysiCell configuration files and rule definitions are provided as Supplementary Files S1-S2 to facilitate full model reproducibility.

### 2. Cell agents and initial conditions

Three agent classes were defined: fibroblasts, macrophages, and capillaries. At simulation start, 100 fibroblasts, 100 macrophages, and 100 capillaries were distributed randomly within the simulation domain. Cell volume was fixed at 2,494 µm³ (nuclear volume 540 µm³) for all agent classes.

Capillaries were implemented as immobile, non-proliferating oxygen-source agents. This formulation imposed a spatially heterogeneous resource landscape that constrained total cell expansion through oxygen availability and local crowding. Unlike analytical ordinary differential equation (ODE) models in which carrying capacity is assigned independently to specific cell types [1, 5], carrying capacity in the present model emerged from shared environmental constraints experienced simultaneously by fibroblasts and macrophages.

Capillary positions remained fixed throughout each simulation to provide reproducible spatial resource geometry. Independent random placement of capillaries across simulations generated distinct spatial terrains sampled across 72 random seeds (Section 6).

Initial cell numbers were selected empirically from pilot simulations. Immediately after initialization, both fibroblast and macrophage populations underwent a transient nucleation phase characterized by partial stochastic extinction before convergence toward stable long-term states. The selected founding population size ensured reliable convergence within the simulation window.

### 3. Oxygen diffusion and shared-resource competition

Oxygen was modeled as a dimensionless proxy for metabolic resources required for cell survival and proliferation. Initial oxygen concentration was set to 10 throughout the simulation domain.

Each capillary secreted oxygen toward a target concentration of 100 at secretion rate 2 /min. Oxygen diffusion coefficient was set to 100,000 µm²/min and decay rate to 10 /min, yielding diffusion lengths on the order of ∼100 µm, comparable to inter-capillary spacing in vascularized tissue. Fibroblasts and macrophages consumed oxygen at identical uptake rates of 100 /min·cell.

This shared-resource formulation differs fundamentally from prior mean-field fibroblast-macrophage circuit models [1, 5], which assume uniform resource access and independently assigned carrying capacities. Here, spatial relationships between cells and capillaries generated emergent carrying-capacity constraints through explicit competition for diffusible resources.

Oxygen parameter values were determined empirically in pilot simulations. Excessively high oxygen supply shifted the dominant constraint from resource competition to purely geometric space filling, thereby obscuring the resource-mediated stochastic failure dynamics studied here. The selected parameter regime preserved oxygen competition as the dominant limiting factor.

Oxygen gradients were recalculated continuously during simulation and contributed to directional migration behavior (Section 5).

### 4. Growth-factor signaling and cell-cycle regulation PDGF and CSF diffusion

Platelet-derived growth factor B (PDGF) and colony-stimulating factor 1 (CSF) were modeled as diffusible dimensionless substrates with diffusion coefficient 5,000 µm²/min and decay rate 5 /min. Initial concentrations were zero. No-flux boundary conditions were applied.

Internalized substrate tracking was enabled for all cells.

Growth-factor-dependent proliferation

Macrophage proliferation was stimulated by CSF using Hill-type activation:

basal proliferation rate: 6.9444 × 10⁻⁵ /min

maximum proliferation rate: 6.9444 × 10⁻⁴ /min half-saturation constant (K₁/₂): 2

Hill coefficient: n = 2

The same functional form was used for PDGF-dependent fibroblast proliferation.

Hill coefficient n = 2 was selected to approximate receptor dimerization behavior of PDGFR and CSF1R signaling systems. Ligand-induced dimerization of PDGFR-β is both necessary and sufficient to drive receptor internalization [9], and CSF-1R undergoes clathrin-dependent endocytosis following ligand binding with subsequent lysosomal degradation [10].

Oxygen-dependent survival

Oxygen reduced necrosis rates in both fibroblasts and macrophages using steep Hill-type survival functions:

maximal necrosis rate: 6.944 × 10⁻⁴ /min

minimal necrosis rate: 6.944 × 10⁻⁵ /min K₁/₂ = 0.1

Hill coefficient: n = 12

The large Hill coefficient implemented a threshold-like transition between viable and necrotic states under oxygen depletion.

Apoptosis rates were fixed at 2.3148 × 10⁻⁴ /min for both cell types.

Growth-factor secretion and enhancer conditions

Fibroblasts constitutively secreted CSF, whereas macrophages secreted PDGF. Secretion rates remained constant across enhancer conditions, while secretion target values were varied to model enhancer-dependent shifts in signaling set points.

Macrophage PDGF secretion target values:

P1 = 1,000

P2 = 2,000

P4 = 4,000

Fibroblast CSF secretion target values:

C1 = 1,000

C2 = 2,000

C4 = 4,000

Enhancer levels were spaced at equal log-scale intervals (factor of 2) to sample multiplicative variation across signaling-strength space, consistent with the log-normal distribution of biological parameters across natural contexts [11].

Fibroblast PDGF secretion was fixed at a level sufficient to sustain homotypic cluster formation and was therefore not treated as enhancer-dependent.

Negative feedback on macrophage PDGF secretion

Macrophages secreted PDGF toward a target density suppressed by CSF signaling. CSF reduced macrophage PDGF secretion toward a minimum target value of 1,000 using a Hill coefficient n = 4 and K₁/₂ = 2.

This negative-feedback branch operated only under P2 and P4 enhancer conditions, where baseline macrophage PDGF secretion exceeded the minimum value. Under P1 conditions, secretion already equaled the minimum and therefore remained unsuppressed.

This asymmetry between homotypic and heterotypic signaling axes was inherited from previous analytical fibroblast-macrophage circuit models [1, 5].

### 5. Cell migration

Fibroblasts and macrophages migrated with identical motility parameters:

migration speed: 1 µm/min

persistence time: 1 min

migration bias: 0.5 Capillaries were immobile.

Migration direction was determined by weighted combinations of normalized substrate gradients. Chemotactic gradients were normalized prior to weighted combination to prevent substrate-specific concentration scales from dominating migration direction.

Fibroblasts migrated up PDGF gradients and macrophages migrated up CSF gradients (weight = 0.7), while both cell types additionally migrated toward oxygen-rich regions (weight = 0.3).

Pilot simulations demonstrated that oxygen-directed migration substantially reduced stochastic circuit failure by facilitating aggregation near capillary-supported regions. Growth-factor-directed migration promoted formation of heterotypic signaling clusters.

Identical motility parameters were assigned to both cell types to isolate the effects of signaling asymmetry from intrinsic motility differences.

### 6. Stochastic terrain sampling

Each enhancer condition was evaluated across 72 independent spatial terrains generated by randomized capillary placement. Simulations were performed on two Apple M3 workstations equipped with an 8-core CPU and 24 GB unified memory. Each simulation required approximately 13 h using 8 OpenMP threads and corresponded to 30,240 simulated minutes (∼3 weeks of biological time). Sequential execution across random seeds 1–72 was automated using a custom Python script (Supplementary File S3). Cell behavior rules, including endocytosis parameter values for each Puptake × Cuptake condition, were configured manually prior to each run series using PhysiCell Studio (version 2.41.7).

Seventy-two terrains were selected to sufficiently sample variability arising from stochastic capillary placement and initial cell distribution while remaining computationally tractable.

Importantly, repeated simulations performed on identical spatial terrains frequently produced divergent outcomes, including coexistence in some runs and stochastic circuit failure in others. This probabilistic divergence indicates that circuit stability cannot be represented solely as a deterministic attractor property of the parameter set.

The simulation design therefore estimated two distinct sources of variability:

1. across-terrain variability arising from capillary geometry
2. within-terrain stochasticity arising from probabilistic cell dynamics

This dual stochastic structure motivated treatment of circuit outcomes as probabilities rather than deterministic states.

### 7. Endocytosis parameters

Endocytosis was modeled as ligand-dependent uptake of PDGF by fibroblasts (Puptake) and CSF by macrophages (Cuptake).

Both uptake processes followed Hill-type activation functions sharing the same K₁/₂ and Hill coefficients used for proliferation signaling (K₁/₂ = 2, n = 2).

For landscape analysis, Puptake and Cuptake were varied independently across parameter grids spanning integer values from 1 to 256 per axis. Each screened value represents the maximum uptake rate for a given condition. The actual uptake rate depends on ligand concentration and is scaled by a Hill activation function (K₁/₂ = 2, n = 2).

The endocytosis-free control condition was defined as Puptake = Cuptake = 0.

Signal-dependent endocytosis was implemented to reflect coupling between ligand occupancy and receptor internalization. This differs from the fixed-rate formulation used in previous analytical models [5].

### 8. Circuit outcome classification

At simulation end (21 days), outcomes were classified into three mutually exclusive categories:

Heterotypic coexistence

At least one macrophage remained associated with a fibroblast cluster.

Homotypic dominance

Fibroblasts persisted whereas macrophages were absent.

Stochastic circuit failure

No fibroblasts remained, regardless of macrophage status.

Fibroblast extinction was selected as the primary failure endpoint because fibroblasts constitute the spatially persistent scaffold of the heterotypic signaling circuit. Pilot time-course analysis indicated that most simulations converged to stable attractors within 1-2 weeks. A 21-day endpoint was selected to ensure convergence across slow trajectories. Representative time-lapse simulations illustrating nucleation, growth, and convergence to heterotypic or homotypic steady states are provided as Supplementary Movie S1.

Outcome probabilities were calculated across 72 spatial terrains:

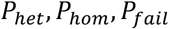

with

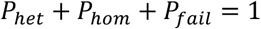

### 9. Composite fitness score and sensitivity analysis

To rank enhancer-endocytosis conditions during screening, a composite fitness score was defined as:

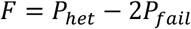

where:

*P*_*het*_= probability of heterotypic coexistence

*P*_*fail*_ = probability of stochastic circuit failure

The composite fitness score 2 was used as an exploratory screening heuristic. Outcome probabilities were subsequently decomposed into *P*_*het*_ and *P*_*fail*_ for mechanistic interpretation independent of the composite weighting scheme.

To evaluate sensitivity to the arbitrary weighting coefficient, ranking analyses were repeated using penalty coefficients ranging from 1 to 4. Relative ordering of high-fitness conditions remained largely preserved across coefficient values, indicating that screening outcomes were determined primarily by the underlying outcome structure rather than by the specific weighting scheme.

Theoretical fitness range:

maximal fitness: +1

minimal fitness: -2

### 10. Pareto-front screening and validation

Endocytosis conditions were identified using a two-stage screening and validation procedure.

Screening stage

All Puptake × Cuptake combinations were initially evaluated using low replication depth (n = 1-2 runs per terrain). Candidate conditions were selected heuristically based on:

1. proximity to the Pareto front in coexistence-failure space
2. exclusion of highly failure-prone conditions
3. inclusion of high-fitness outliers

Pareto-optimal conditions were defined as parameter combinations for which no alternative condition simultaneously increased coexistence fraction while reducing stochastic circuit failure.

Because Pareto-front structure remained uncertain under limited replication depth, the screening criteria were defined heuristically prior to validation analysis.

Validation stage

Selected conditions (four per enhancer combination) were re-evaluated independently using: 72 terrains

8 independent stochastic replicates per terrain

576 simulations per condition

Screening-stage simulations were excluded from validation analysis to avoid winner’s-curse bias.

Statistical analysis

Differences among endocytosis conditions were initially evaluated using Kruskal-Wallis tests; all comparisons passed the omnibus test prior to post-hoc analysis.

Post-hoc comparisons were structured as directional comparisons against matched endocytosis-free controls using one-tailed Mann-Whitney U tests with Bonferroni correction (k = 4 comparisons per enhancer condition).

One-tailed testing was selected on the basis of a directional a priori hypothesis: previous analytical studies of fibroblast-macrophage circuits demonstrated that receptor-mediated endocytosis stabilizes communicating cell systems by suppressing growth-factor accumulation [5], predicting that endocytosis-enabled conditions should exhibit lower failure rates and higher fitness than endocytosis-free controls. Although the present spatial agent-based framework differs fundamentally from the deterministic ODE formulation of Adler et al. in its treatment of spatial heterogeneity and shared resource competition, the predicted direction of endocytosis-dependent stabilization is shared across both frameworks, justifying a directional test.

Statistical analyses were performed using SPSS version 31 (IBM Corp., Armonk, NY, USA).

### 11. Hypothetical homotypic baseline simulations

To estimate failure risk associated with asymmetric CSF dependence, we simulated a hypothetical homotypic baseline architecture in which both cell populations produced and responded exclusively to PDGF.

In this configuration:

CSF production = 0

macrophage CSF response = absent

both populations responded to PDGF identically

Migration followed combined PDGF and oxygen gradients for both cell types. Initial conditions and spatial terrain sampling were identical to those used in heterotypic simulations.

In the homotypic baseline, two failure criteria were evaluated: simultaneous extinction of both cell populations (A and B), and extinction of either cell population (A or B). The A and B criterion corresponds most directly to the fibroblast extinction endpoint used in the heterotypic circuit, in which circuit failure was defined by loss of fibroblasts regardless of macrophage status. The A or B criterion was included as a supplementary measure to assess partial population collapse.

Differences among groups were assessed using a Kruskal-Wallis test followed by post-hoc two-tailed Mann-Whitney U tests with Bonferroni correction (k = 2 comparisons against the heterotypic control). The A or B criterion showed a trend toward lower failure frequency compared to the heterotypic circuit but did not reach significance after correction (p = 0.045, uncorrected), and is reported descriptively.

Homotypic baseline simulations were performed under P1P1 enhancer conditions to provide a conservative estimate of failure risk in the absence of CSF dependence. This condition was selected to minimize autocrine signaling strength, thereby isolating the structural effect of circuit asymmetry from enhancer-dependent differences in single-population viability.

## Results

Asymmetric CSF dependence increases stochastic circuit failure relative to a hypothetical homotypic baseline

To estimate the systems-level cost associated with asymmetric CSF dependence, we compared the fibroblast-macrophage heterotypic circuit against a hypothetical homotypic baseline architecture in which both cell populations produced and responded exclusively to PDGF (Figure 1A).

**Figure 1.**
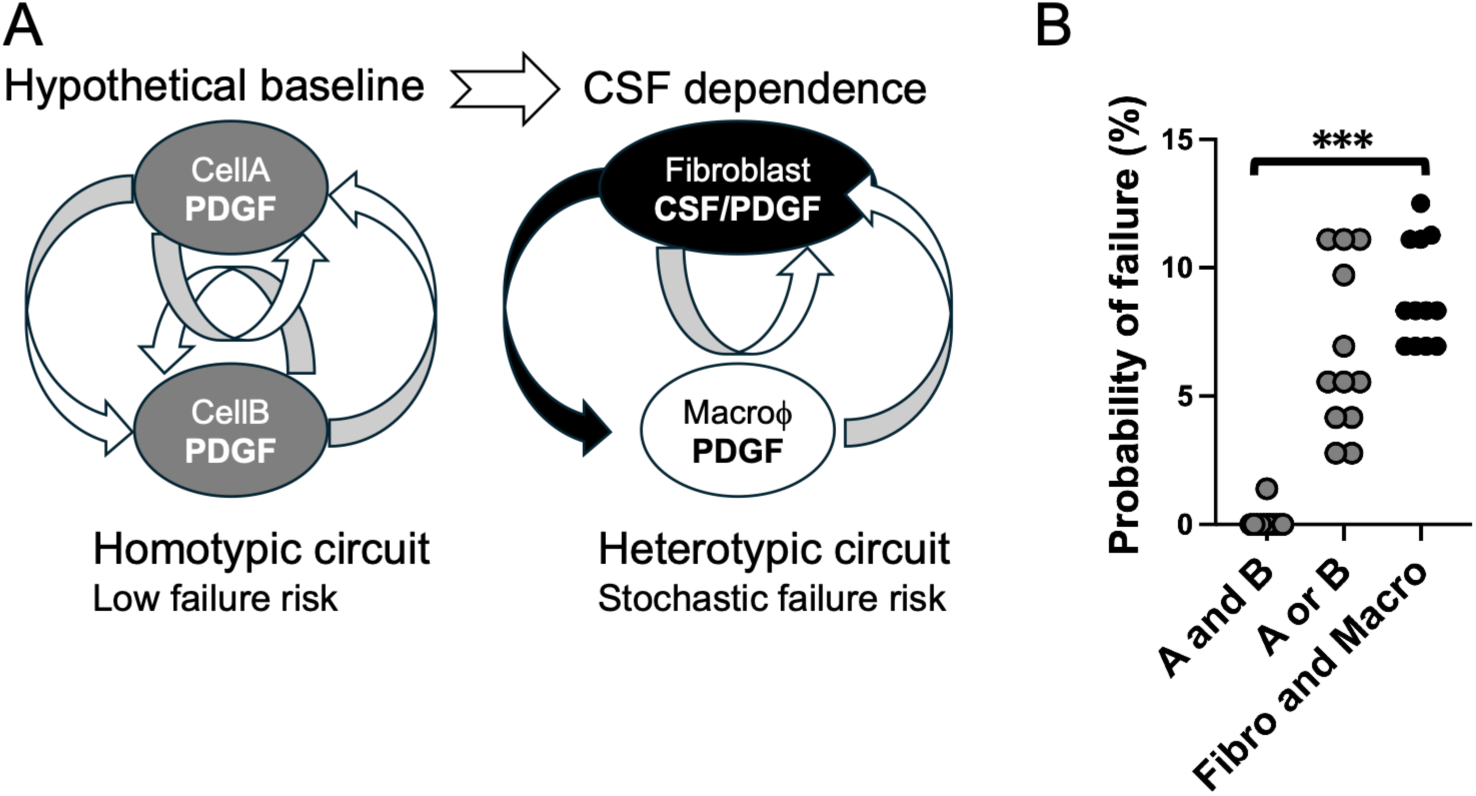
Introduction of asymmetric CSF dependence increases stochastic circuit failure relative to a hypothetical homotypic baseline. (A) Schematic comparison between a hypothetical homotypic baseline architecture and the fibroblast-macrophage heterotypic circuit. In the homotypic baseline, both cell types produce and respond to PDGF. In the heterotypic circuit, fibroblasts produce CSF to support macrophages, whereas macrophages produce PDGF to support fibroblasts, generating asymmetric trophic dependence. (B) Comparison of failure fraction among the hypothetical homotypic baseline circuit (A and B: simultaneous extinction of both cell types; A or B: extinction of either cell type) and the heterotypic fibroblast-macrophage circuit (Fibro and Macro: fibroblast and macrophage extinction) under endocytosis-free conditions (P1C1). Each dot represents one simulation run using the same spatial seeds (n = 12 runs × 72 spatial configurations). Horizontal lines indicate means. Differences among groups were assessed using a Kruskal-Wallis test (omnibus test passed) followed by post-hoc two-tailed Mann-Whitney U tests with Bonferroni correction (k = 2): *** p < 0.001 (A and B vs. Fibro and Macro). A or B extinction also occurred at lower frequency than heterotypic circuit failure, but did not reach significance after Bonferroni correction (p = 0.045, uncorrected).

Under endocytosis-free conditions, the heterotypic circuit exhibited substantially higher stochastic circuit failure fraction than the homotypic baseline architecture (*P* < 0.001, *n* = 12 runs × 72 spatial configurations, Mann-Whitney *U* test, two-tailed, Figure 1B). In the homotypic baseline, extinction required simultaneous loss of both populations, whereas the heterotypic circuit became vulnerable to fibroblast extinction through indirect macrophage-mediated resource depletion.

The A or B criterion, which counts extinction of either population regardless of the other, did not reach significance after Bonferroni correction (p = 0.045, uncorrected), consistent with the expectation that partial population loss is less catastrophic in a homotypic circuit where the surviving population maintains signaling autonomy. The primary comparison therefore focused on simultaneous extinction of both populations (A and B), which most directly parallels the functional definition of circuit failure in the heterotypic system.

This result indicates that introduction of asymmetric CSF dependence increases susceptibility to stochastic collapse under shared-resource competition. Importantly, however, subsequent analyses demonstrated that this vulnerability could be selectively suppressed by endocytosis-dependent negative feedback.

Endocytosis suppresses stochastic circuit failure across enhancer conditions

To investigate how endocytosis modulates circuit stability, we screened independent combinations of fibroblast PDGF uptake (Puptake) and macrophage CSF uptake (Cuptake) across nine enhancer conditions spanning fourfold variation in PDGF and CSF secretion targets (P1-P4 × C1-C4).

The stochastic nature of circuit dynamics is illustrated in Supplementary Movie S1, which shows representative simulations under heterotypic coexistence. Pareto-front analysis consistently identified endocytosis-enabled conditions displaced toward lower stochastic circuit failure fractions relative to endocytosis-free controls (Figure 2). Although coexistence remained broadly achievable across many parameter combinations, low-failure conditions clustered preferentially in regions of elevated Cuptake.

**Figure 2.**
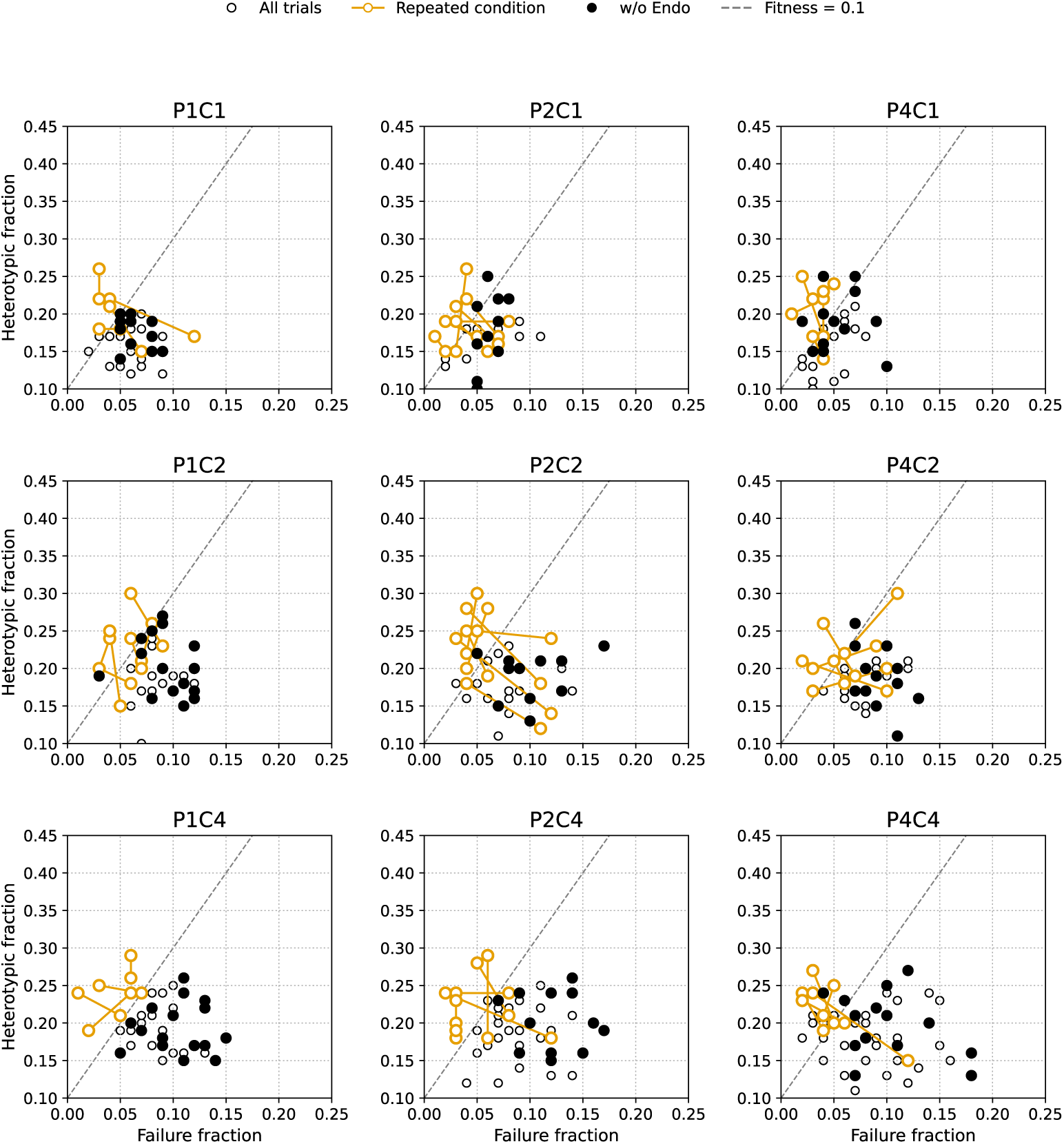
Pareto-front screening of endocytosis parameters across nine enhancer conditions. Puptake denotes PDGF endocytosis by fibroblasts, and Cuptake denotes CSF endocytosis by macrophages. Each panel shows heterotypic fraction versus failure fraction for all screened Puptake × Cuptake parameter combinations under a given enhancer condition (P1-P4 × C1-C4; PDGF enhancer × CSF enhancer). Each replicate was evaluated using 72 independent spatial seeds. Open circles indicate all screened parameter combinations. Orange connected points indicate Pareto-optimal conditions, defined as parameter combinations for which no alternative condition simultaneously improved heterotypic fraction and reduced failure fraction. Filled black circles indicate endocytosis-free controls (w/o Endo). The dashed line denotes Fitness = 0.1, where Fitness is defined as *F* = *P*_*het*_ − 2*P*_*fail*_. Pareto-optimal conditions were consistently displaced toward reduced failure fraction relative to endocytosis-free controls across multiple enhancer conditions.

To validate these trends independently from the exploratory screening stage, selected parameter combinations were re-evaluated using 72 spatial terrains and 8 stochastic replicates per terrain (576 simulations per condition). Statistical comparison against matched endocytosis-free controls demonstrated that multiple endocytosis-enabled conditions significantly improved overall fitness (Figure 3).

**Figure 3.**
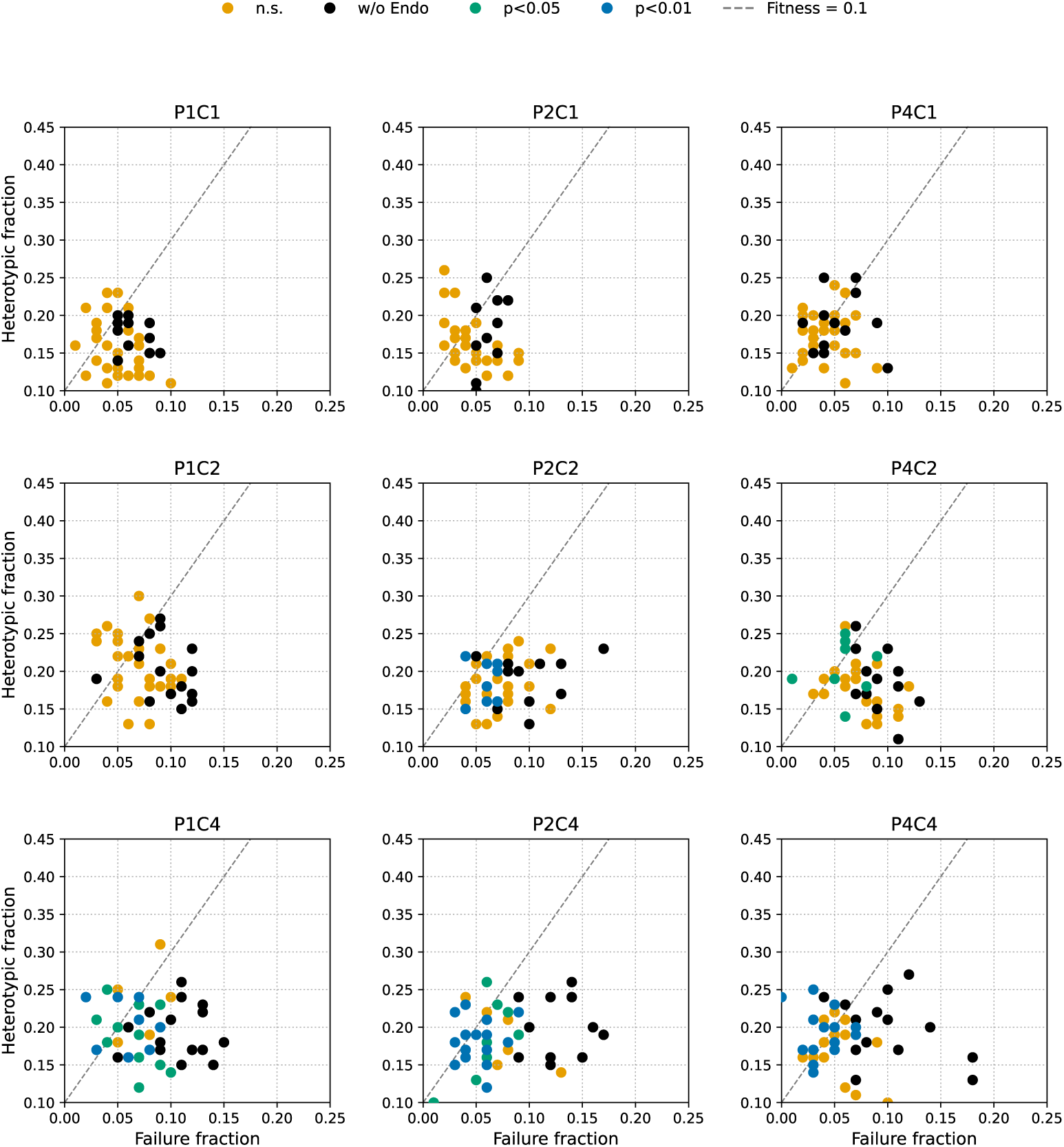
Validation analysis confirms endocytosis-dependent suppression of stochastic circuit failure. Validation dataset excluding screening-stage runs (n = 8 independent replicates × 72 spatial seeds per condition). Each dot represents one validated Puptake × Cuptake parameter combination plotted in heterotypic fraction versus failure fraction space. Color indicates statistical comparison against the matched endocytosis-free control within each enhancer condition using one-tailed Mann-Whitney U tests with Bonferroni correction for four comparisons: orange, not significant; teal, p < 0.05; blue, p < 0.01. Filled black circles indicate endocytosis-free controls. The dashed line denotes Fitness = 0.1. Significant conditions clustered preferentially at low failure fractions, indicating that endocytosis primarily stabilized the circuit through suppression of stochastic failure.

Notably, significant conditions localized preferentially in low-failure regions rather than in regions with unusually elevated coexistence fraction. This suggested that endocytosis-dependent fitness improvement arose primarily through suppression of stochastic circuit failure rather than enhancement of coexistence probability itself.

To interpret the pattern of significant and non-significant endocytosis effects across enhancer conditions, we examined baseline fitness variation among endocytosis-free controls. Composite fitness scores varied significantly across the nine enhancer combinations (Kruskal-Wallis P < 0.001). Post-hoc pairwise comparisons revealed that conditions with elevated CSF enhancer strength (P1C4, P2C4) exhibited significantly lower baseline fitness than P4C1 (adjusted P = 0.017 and 0.002, respectively), with P2C2 also differing significantly from P4C1 (adjusted P = 0.048). In contrast, all three PxC1 conditions showed comparable baseline fitness with no significant pairwise differences among them (all adjusted P > 0.05), indicating that the low-failure, high-fitness regime was a structural feature of low CSF enhancer conditions rather than a property specific to P4C1. This pattern is consistent with a floor effect across PxC1 conditions: the limited dynamic range of baseline failure rates under low CSF signaling constrained the statistical detectability of endocytosis-dependent stabilization, with the exception of P2C1, where endocytosis-dependent stabilization remained statistically detectable (Figure 7).

Composite fitness improvement is driven primarily by reduced failure fraction

To determine which outcome probabilities contributed to fitness improvement, we decomposed the composite fitness metric into its constituent coexistence and failure fractions.

Across enhancer conditions, coexistence probability remained relatively stable despite large variation in endocytosis parameters (Figures 4). Kruskal-Wallis tests confirmed that endocytic activity did not significantly alter heterotypic fraction in any of the nine enhancer conditions (all *P* > 0.05), indicating that endocytosis acts selectively on circuit failure rather than on the establishment or maintenance of heterotypic coexistence. In contrast, stochastic circuit failure exhibited pronounced sensitivity to Cuptake (Figures 6 and 7). Significant reductions in failure fraction extended beyond conditions exhibiting statistically significant fitness improvement, indicating that failure suppression was more sensitive to endocytosis-dependent stabilization than the composite fitness metric itself.

**Figure 4.**
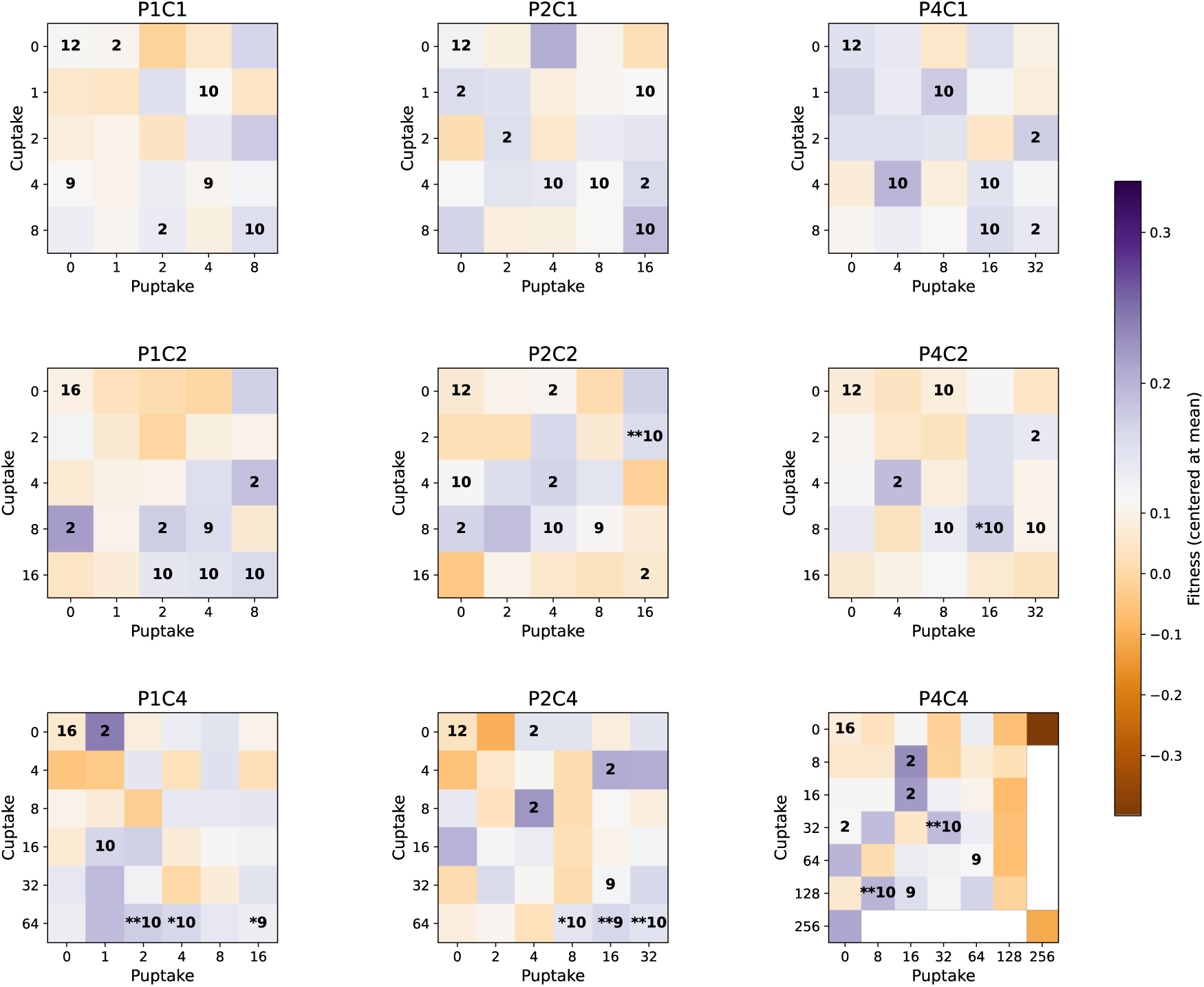
Fitness landscapes across Puptake × Cuptake parameter space. Heatmaps display mean Fitness values as a function of Puptake (x-axis) and Cuptake (y-axis) for each enhancer condition (P1-P4 × C1-C4). Fitness was defined as *F* = *P*_*het*_ − 2*P*_*fail*_. Values are displayed as deviations from the mean fitness among the entire data set to emphasize relative landscape structure. A diverging PuOr colormap was applied; orange indicates above-mean fitness, whereas purple indicates below-mean fitness, with white corresponding to zero deviation. Numbers within each cell indicate the number of validation replicates. Asterisks denote statistical significance relative to the matched endocytosis-free control using one-tailed Mann-Whitney U tests with Bonferroni correction (directional hypothesis based on Adler et al. [5]): * p < 0.05; ** p < 0.01. Elevated fitness regions emerged preferentially at intermediate-to-high Cuptake values.

**Figure 5.**
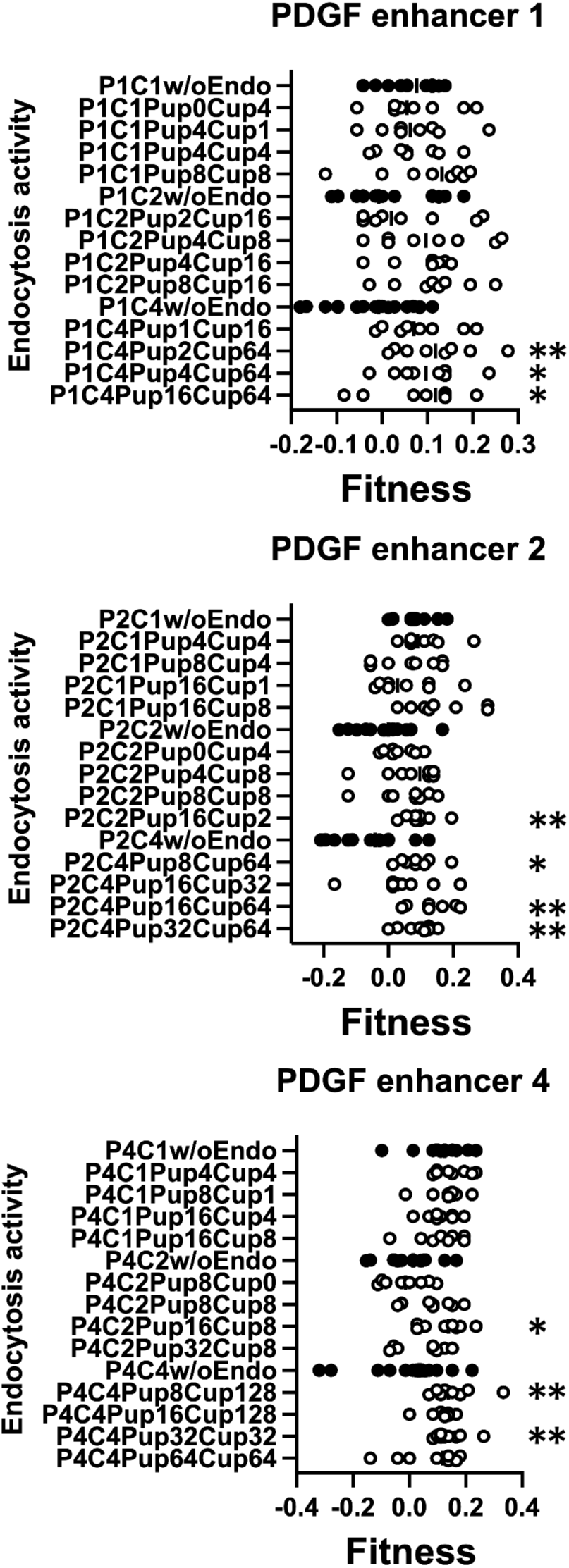
Distribution of fitness values across validated endocytosis parameter combinations. Dot plots showing Fitness values for validated Puptake × Cuptake parameter combinations selected from Pareto-front validation. Filled circles indicate endocytosis-free controls. Rows correspond to PDGF enhancer conditions (P1, P2, P4), and columns correspond to CSF enhancer conditions (C1, C2, C4). Each dot represents one independent simulation replicate. Asterisks indicate statistically significant improvement relative to the matched endocytosis-free control using one-tailed Mann-Whitney U tests with Bonferroni correction (directional hypothesis based on Adler et al. [5]): * p < 0.05; ** p < 0.01. Multiple distinct Puptake × Cuptake combinations yielded significant fitness improvement across enhancer conditions.

**Figure 6.**
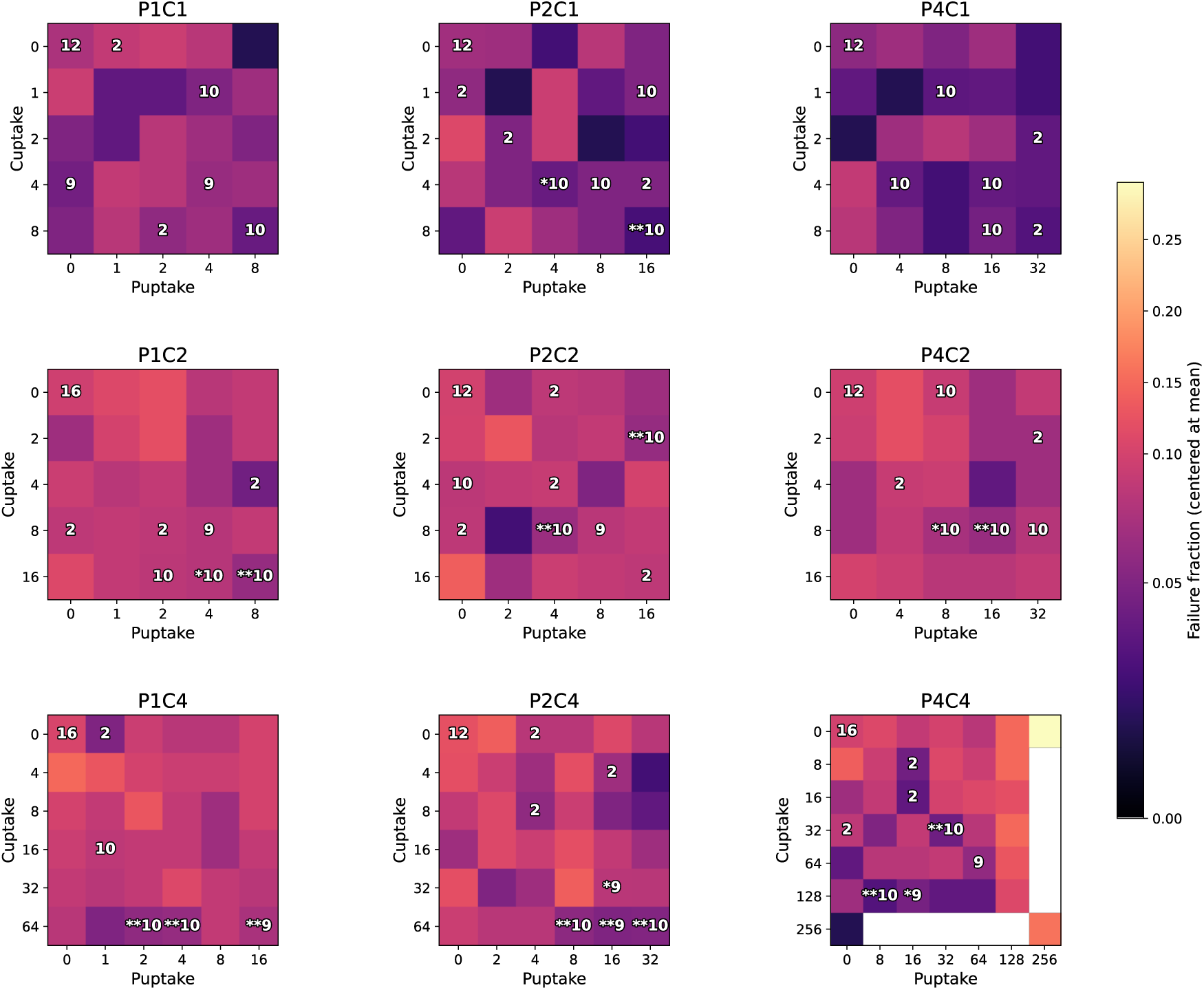
Failure landscapes identify CSF endocytosis as the dominant stabilizing parameter. Heatmaps display mean failure fraction as a function of Puptake (x-axis) and Cuptake (y-axis) for each enhancer condition (P1-P4 × C1-C4). Values are displayed as deviations from the mean failure fraction among the entire data set to emphasize relative landscape structure. A magma colormap was applied; lighter colors indicate above-mean failure fraction, whereas darker colors indicate below-mean failure fraction. Numbers within each cell indicate the number of validation replicates. Asterisks denote statistical significance relative to the matched endocytosis-free control using one-tailed Mann-Whitney U tests with Bonferroni correction (directional hypothesis based on Adler et al. [5]): * p < 0.05; ** p < 0.01. Significant reductions in failure fraction extended beyond conditions showing significant improvement in the composite fitness metric, indicating that stochastic failure suppression is more sensitive to endocytic stabilization than overall fitness. Low-failure regions were concentrated at elevated Cuptake values across multiple enhancer conditions.

**Figure 7.**
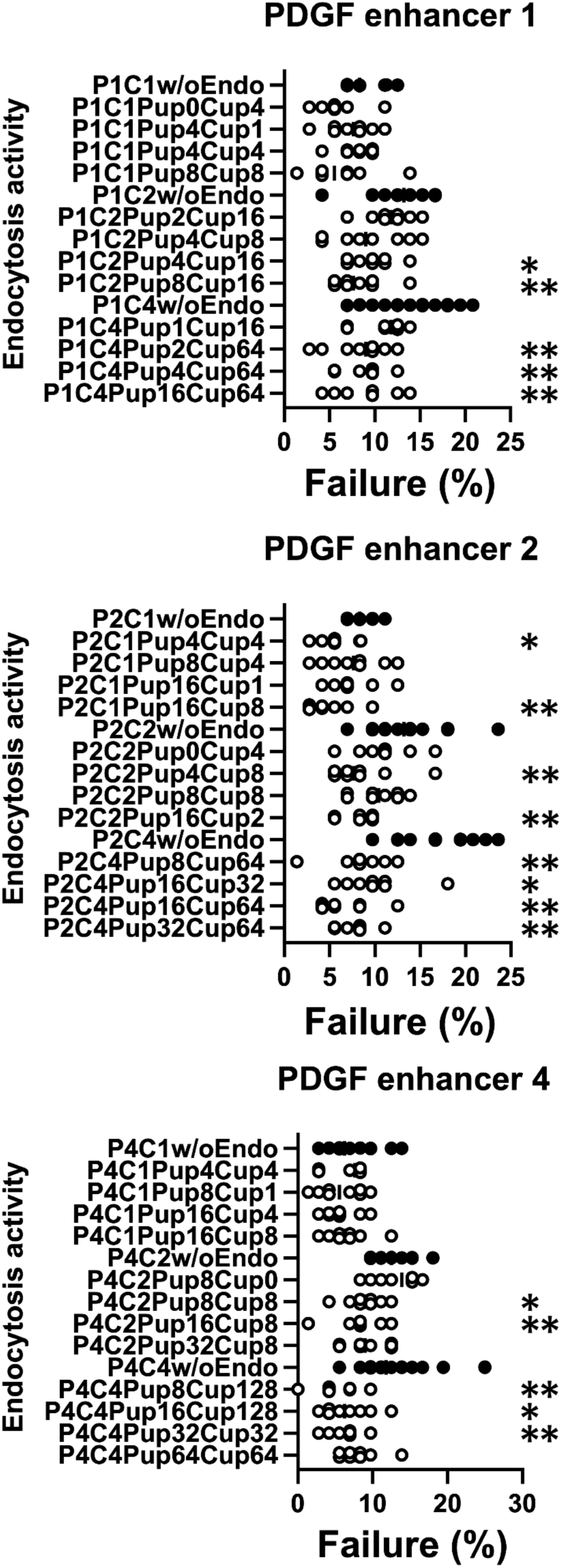
Endocytosis reduces stochastic circuit failure across multiple enhancer conditions. Dot plots showing failure fraction for validated Puptake × Cuptake parameter combinations selected from Pareto-front validation. Filled circles indicate endocytosis-free controls. Rows correspond to PDGF enhancer conditions (P1, P2, P4), and columns correspond to CSF enhancer conditions (C1, C2, C4). Each dot represents one independent simulation replicate. Asterisks indicate statistically significant reduction in failure fraction relative to the matched endocytosis-free control using one-tailed Mann-Whitney U tests with Bonferroni correction (directional hypothesis based on Adler et al. [5]): * p < 0.05; ** p < 0.01. Significant reductions in failure fraction were preferentially associated with elevated Cuptake values, whereas broad ranges of Puptake values yielded comparable stabilization under high-Cuptake conditions.

These results demonstrated that endocytosis did not substantially increase the probability of establishing heterotypic coexistence once stable clusters formed. Instead, endocytosis acted primarily by preventing catastrophic extinction events that otherwise terminated the circuit before long-term coexistence could emerge.

Accordingly, mechanistic interpretation of the simulations shifted from “coexistence enhancement” toward “stochastic failure suppression.”

Spatial resource competition identifies CSF uptake as the dominant stabilizing feedback

Fitness landscapes across Puptake × Cuptake parameter space revealed strong asymmetry between the PDGF and CSF uptake axes (Figure 4). Elevated-fitness regions emerged preferentially at intermediate-to-high Cuptake values across multiple enhancer conditions, whereas broad variation in Puptake frequently produced relatively similar outcomes.

This asymmetry became substantially clearer in the failure landscapes (Figure 6). Low-failure regions were concentrated almost exclusively at elevated Cuptake values, whereas insufficient Cuptake consistently failed to suppress collapse regardless of Puptake level.

Importantly, this systems-level asymmetry closely parallels the stabilizing principle proposed in analytical fibroblast-macrophage circuit models [1, 5]. In those studies, fibroblasts were treated as near carrying capacity whereas macrophages remained far from carrying capacity, predicting that stabilization should depend primarily on negative feedback acting on the macrophage-supporting CSF axis.

In contrast, the present model imposed no independent carrying capacities. Instead, fibroblasts and macrophages competed explicitly for shared oxygen and space. Under this formulation, excessive CSF signaling drove macrophage overexpansion, depletion of shared resources, and eventual fibroblast extinction.

Despite these fundamentally different assumptions, both analytical and spatial frameworks converged on the same systems-level conclusion: stabilization of the macrophage-supporting CSF axis is substantially more important than stabilization of the PDGF axis.

Permissive stabilization landscapes emerge along the PDGF uptake axis

Failure landscapes revealed a pronounced asymmetry in parameter sensitivity along the Puptake axis. Under high-Cuptake conditions, broad ranges of Puptake values yielded similarly low stochastic circuit failure fractions (Figure 6 and 7, P1C2, P4C2, P1C4, P2C4 and P4C4). This permissive landscape was especially prominent under high-CSF enhancer conditions (C4), where elevated Cuptake robustly suppressed collapse across diverse Puptake values.

Once Cuptake exceeded threshold levels, multiple distinct Puptake values produced comparable stabilization outcomes. In contrast, low Cuptake conditions remained failure-prone regardless of Puptake magnitude. One exception was the P2C2 condition at Puptake16/Cuptake2, which showed reduced failure despite low Cuptake (Figure 6 and 7).

This asymmetry indicates that Cuptake functions as the dominant stabilizing feedback in the circuit, whereas Puptake primarily modulates non-catastrophic competitive balance. Mechanistically, excess PDGF favored fibroblast-dominant outcomes without necessarily eliminating the circuit, whereas excess CSF promoted macrophage overexpansion capable of exhausting shared resources and driving irreversible fibroblast extinction.

Thus, the stabilizing landscape was threshold-sensitive along the CSF uptake axis but broadly permissive along the PDGF uptake axis, generating a permissive parameter regime in which robust stabilization emerged without requiring precise tuning of PDGF endocytosis.

Composite fitness rankings are robust to weighting-coefficient variation

The composite fitness score used during screening incorporated a penalty coefficient weighting stochastic circuit failure more strongly than coexistence loss. To determine whether screening outcomes depended critically on this arbitrary coefficient choice, we repeated ranking analyses using alternative penalty coefficients ranging from 1 to 4.

Relative ordering of high-fitness parameter combinations remained largely preserved across coefficient values (Supplementary Figure 1). This robustness indicates that the dominant structure of the fitness landscape was determined primarily by the underlying outcome probabilities rather than by the precise numerical weighting scheme.

Importantly, subsequent decomposition of the fitness metric demonstrated that endocytosis-dependent stabilization emerged almost entirely through reduced stochastic circuit failure rather than altered coexistence probability. Thus, the principal biological conclusions of the study remained independent of the specific composite weighting formulation used during exploratory screening.

## Discussion

### Endocytosis suppresses stochastic extinction rather than promoting coexistence

The principal finding of this study is that endocytic activity selectively suppresses stochastic circuit failure without measurably increasing heterotypic coexistence. Endocytosis-active parameter combinations significantly reduced failure fraction in all enhancer conditions except P4C1 (Figure 6 and 7), whereas heterotypic fraction remained statistically unchanged relative to endocytosis-free controls. This functional dissociation indicates that endocytosis does not actively promote coexistence once fibroblast-macrophage clusters are established; rather, it reduces the probability that the circuit collapses before a stable coexistence state can emerge.

This interpretation extends prior theoretical work on fibroblast-macrophage circuit stability. Zhou et al. proposed that stable two-cell systems require a “spring-and-ceiling” architecture in which one cell type is constrained by carrying capacity while the other remains proliferative, thereby necessitating negative feedback on the corresponding trophic factor [1]. Adler et al. subsequently demonstrated analytically that receptor-mediated endocytosis can stabilize such communicating circuits by providing robust negative feedback on growth-factor accumulation [5]. In those formulations, endocytosis stabilizes a deterministic ON-state equilibrium. Our results extend this principle into a spatial and stochastic regime: under shared oxygen and space constraints, endocytosis acts primarily by suppressing catastrophic extinction events rather than by enhancing coexistence itself.

### Shared-resource competition spatially reinterprets the carrying-capacity framework

The present framework differs fundamentally from the carrying-capacity assumptions used in the analytical models of Zhou et al. and Adler et al. In those studies, fibroblasts were experimentally observed to undergo density-dependent growth arrest in culture, whereas macrophages remained proliferative, motivating a formulation in which fibroblasts were treated as near carrying capacity and macrophages as far from carrying capacity [1]. However, this distinction emerged under culture conditions in which fibroblasts formed relatively stationary adherent populations, whereas macrophages exhibited highly motile behavior.

In our spatial framework, carrying capacity was not assigned independently to a single cell type. Instead, both fibroblasts and macrophages competed explicitly for shared oxygen and physical space, allowing carrying capacity to emerge from common environmental constraints. Under this formulation, excessive CSF-driven macrophage expansion depleted shared resources and eliminated fibroblasts through competitive exclusion, thereby generating a stochastic failure mode absent from independent-capacity formulations.

Importantly, despite these fundamentally different assumptions, both frameworks converged on the same systems-level conclusion: stabilization of the macrophage-supporting CSF axis is substantially more critical than stabilization of the PDGF axis. In our simulations, Cuptake exhibited a strict threshold requirement for failure suppression, whereas Puptake remained broadly permissive across a wide parameter range. This convergence suggests that asymmetric sensitivity to CSF accumulation may represent a robust design principle of interdependent fibroblast-macrophage circuits rather than an artifact of a specific modeling formalism.

### Asymmetric parameter sensitivity identifies CSF endocytosis as the critical stabilizing parameter

The failure landscape exhibited a pronounced sloppy topology along the Puptake axis. In C4 enhancer conditions, broad ranges of Puptake values yielded nearly equivalent reductions in failure fraction provided that Cuptake exceeded a threshold level. By contrast, low Cuptake conditions consistently failed to suppress collapse regardless of Puptake value. This asymmetry identifies CSF endocytosis as the rate-limiting stabilizing feedback in the circuit.

The concept of sloppy parameter sensitivity was originally introduced in systems biology models to describe systems in which behavior is highly sensitive to a limited subset of parameter combinations while remaining broadly insensitive to variation along other directions [12, 13]. The failure landscape observed here shares this structural feature along the Puptake axis: once Cuptake exceeded a threshold level, broad variation in Puptake produced comparable stabilization outcomes, generating a permissive parameter regime that did not require precise tuning of PDGF endocytosis.

Mechanistically, this asymmetry reflects the unequal consequences of ligand excess. Excess PDGF primarily accelerates fibroblast proliferation and favors homotypic dominance, whereas excess CSF produces macrophage overexpansion that consumes shared oxygen and space, ultimately driving fibroblast extinction and irreversible circuit collapse. Endocytosis therefore functions asymmetrically within the circuit architecture: Cuptake acts as a critical brake preventing catastrophic resource monopolization, whereas Puptake primarily modulates non-catastrophic competitive balance.

### Endocytosis may function as a robustness architecture against collapse-prone mutualistic dynamics

More broadly, the observed extinction-suppressing effect of endocytosis resembles stabilizing mechanisms described in mutualistic network theory, in which gradual parameter variation can produce nonlinear transitions once critical thresholds are crossed. Our results extend this concept to spatially embedded multicellular signaling circuits under shared-resource competition.

Notably, the accelerator-threshold behavior observed across C1-C4 enhancer conditions is consistent with this interpretation. Under low CSF enhancer conditions, macrophage expansion remained insufficient to drive frequent collapse, leaving limited room for measurable endocytosis-dependent stabilization. In contrast, high CSF enhancer conditions entered a collapse-prone regime in which Cuptake-mediated negative feedback robustly suppressed extinction. The emergence of permissive stabilization landscapes specifically within these collapse-prone regimes suggests that endocytosis may widen the viable parameter space of interdependent circuits without requiring finely tuned enhancer activity.

This interpretation is also consistent with broader principles of robustness in biological networks, in which functional architectures evolve to maintain system stability despite substantial parameter variation [14]. Although the present study does not explicitly simulate adaptive evolution, the convergence between analytical homeostasis models and spatial stochastic simulations suggests that CSF-focused negative feedback may represent a robust systems-level stabilization architecture compatible with interdependent fibroblast-macrophage systems.

The collapse dynamics observed here share structural features with mutualistic network theory, in which mutually dependent populations can abruptly collapse once stabilizing constraints fail [15]. In both cases, nonlinear feedback and shared resource constraints define a collapse threshold beyond which stochastic perturbations drive irreversible extinction. The present results suggest that endocytic negative feedback may function as a stabilizing constraint analogous to those that prevent catastrophic transitions in ecological mutualistic systems.

### Limitations

The present framework does not explicitly simulate adaptive evolutionary processes such as mutation, selection, or heritable parameter change. Accordingly, the evolutionary interpretations presented here should be understood as robustness-based plausibility arguments rather than direct reconstructions of circuit evolution.

In addition, enhancer strengths, endocytosis rates, and oxygen-dependent resource competition were modeled as abstract control parameters rather than experimentally calibrated molecular rate constants or tissue-specific physiological constraints. Oxygen was implemented as a generic diffusible proxy for shared environmental limitation, and the observed collapse dynamics are therefore expected to generalize to other shared constraints including nutrients, ECM-limited space, and diffusible metabolic support.

Despite these simplifications, the convergence between analytically derived stabilization principles and spatially emergent failure suppression suggests that the observed asymmetry of the CSF axis reflects a systems-level property that is robust across distinct modeling assumptions.

## Author Contributions

K.I. conceived and designed the computational experiments, performed the majority of simulations, conducted all formal statistical analyses, interpreted the results, created all visualizations, and wrote the original draft of the manuscript.

Y.I. implemented the PhysiCell and PhysiCell Studio model configurations, developed the automated simulation script for 72 random seed execution, and contributed to simulation runs.

M.H. originated the conceptual framework of the study, contributed to PhysiCell and PhysiCell Studio implementation, and provided critical review of the manuscript.

All authors reviewed and approved the final manuscript.

## Data Availability

All simulation configuration files and cell behavior rules required to reproduce the agent-based model are provided as Supplementary Files S1 (PhysiCell_settings.xml) and S2 (detailed_rules.txt). Raw simulation outcome data underlying all figures and statistical analyses are provided as Supplementary Tables S1 and S2. Complete simulation output archives are stored locally and are available from the corresponding author upon reasonable request.

## Acknowledgement

This work was supported by JSPS KAKENHI (grant number 24K15733).

## Supplementary data

**Supplementary Figure S1.**
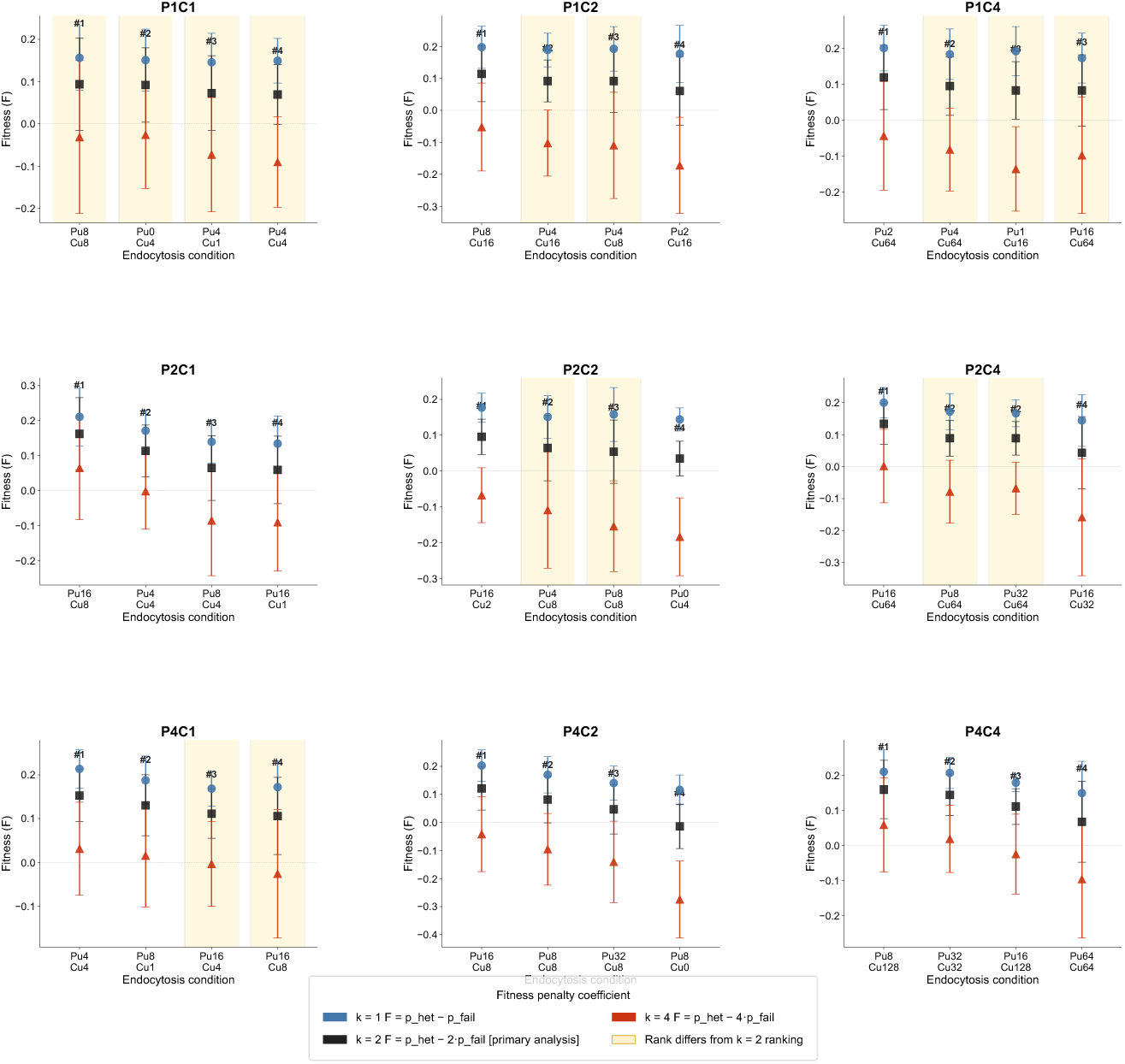
Sensitivity of Pareto condition ranking to fitness penalty coefficient (k) Mean fitness (F) of the four endocytosis conditions selected per enhancer combination, evaluated under three penalty coefficients: k = 1 (*F* = *P*_*het*_ – *P*_*fail*_; blue circles), k = 2 (*F* = *P*_*het*_ − 2*P*_*fail*_; black squares; primary analysis), and k = 4 (*F* = *P*_*het*_ − 4*P*_*fail*_; red triangles). Each panel corresponds to one of the nine enhancer combinations (P1C1 through P4C4). The x-axis shows the four endocytosis conditions sorted in ascending order of k = 2 rank (rank #1 = highest fitness under primary analysis); bold rank labels (#1-#4) indicate k = 2 rankings. Error bars represent standard deviation (SD) across simulation runs (n = 8 runs × 72 spatial configurations per condition). Yellow shading marks conditions whose rank differs between k values. The dotted horizontal line indicates F = 0. Note that all three fitness functions yield negative mean F values under k = 4 for most conditions, reflecting the amplified penalty on failure probability; this does not alter the relative ranking of conditions within each panel. The top-ranked condition (rank #1 under k = 2) was preserved under k = 1 and k = 4 in 8 of 9 enhancer combinations; in P1C1, ranks #1 and #2 were exchangeable across k values owing to nearly identical mean fitness scores (ΔF < 0.005). These results indicate that the identity of Pareto-dominant conditions is robust to the choice of penalty coefficient, supporting the validity of the primary analysis (k = 2).

**Supplementary Figure S2.**
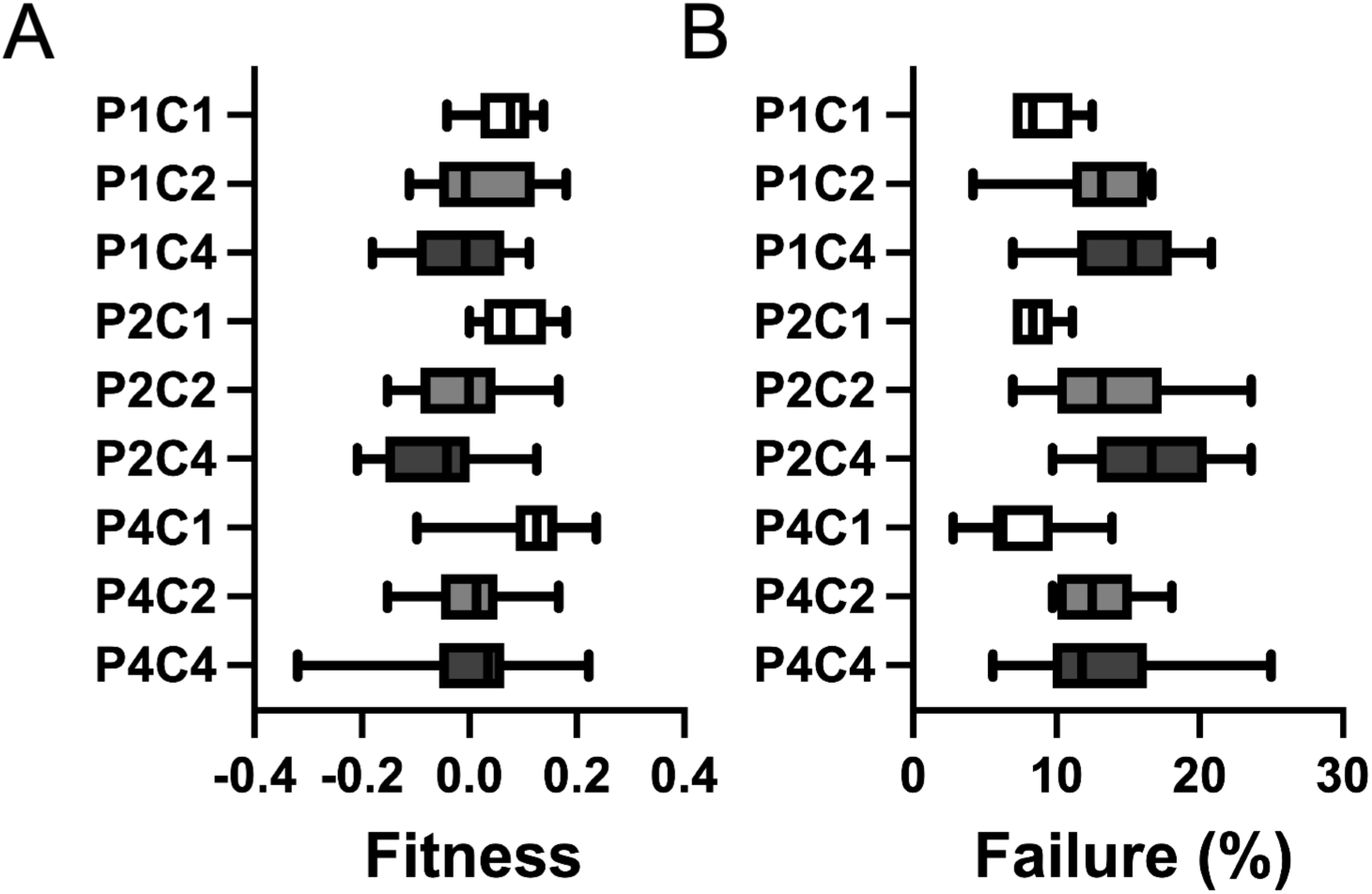
Baseline fitness and failure rates vary with CSF enhancer strength under endocytosis-free conditions. Box plots showing the distribution of composite fitness scores (A) and stochastic circuit failure rates (B) across nine enhancer conditions under endocytosis-free conditions (Puptake = Cuptake = 0). Each condition is represented by all endocytosis-free control runs from the validation dataset (n = 72 spatial terrains per condition). Box fill indicates CSF enhancer level: white, PxC1; light gray, PxC2; dark gray, PxC4. Boxes indicate interquartile range; horizontal lines within boxes indicate medians; whiskers extend to the most extreme values within 1.5 × IQR. Both fitness and failure rates varied significantly across enhancer combinations (Kruskal-Wallis P < 0.001 for both). Post-hoc pairwise comparisons (Mann-Whitney U, Bonferroni correction) identified significantly lower baseline fitness in P1C4 and P2C4 relative to P4C1 (adjusted P = 0.017 and P = 0.002, respectively), with P2C2 also differing significantly from P4C1 (adjusted P = 0.048). Within each PDGF enhancer level, failure rates and fitness scores increased and decreased monotonically with CSF enhancer strength, respectively (PxC1 < PxC2 < PxC4 for failure; PxC1 > PxC2 > PxC4 for fitness), consistent with a structural floor effect under low CSF signaling conditions. Among all nine enhancer combinations, P4C1 exhibited the lowest median failure rate, indicating that high PDGF enhancer strength combined with low CSF enhancer strength produced the most stochastically stable baseline circuit state.

**Supplementary Table S1.**
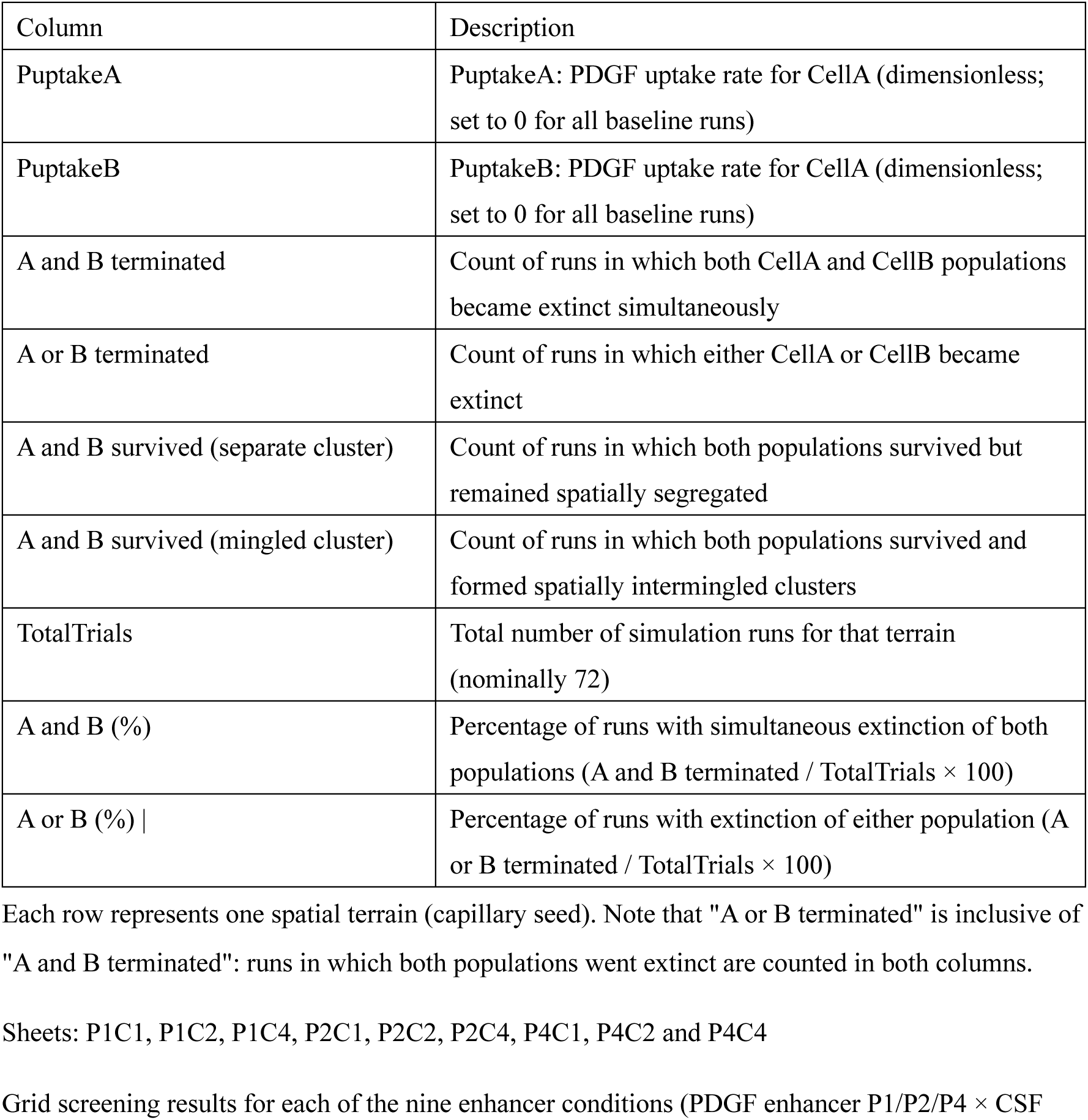
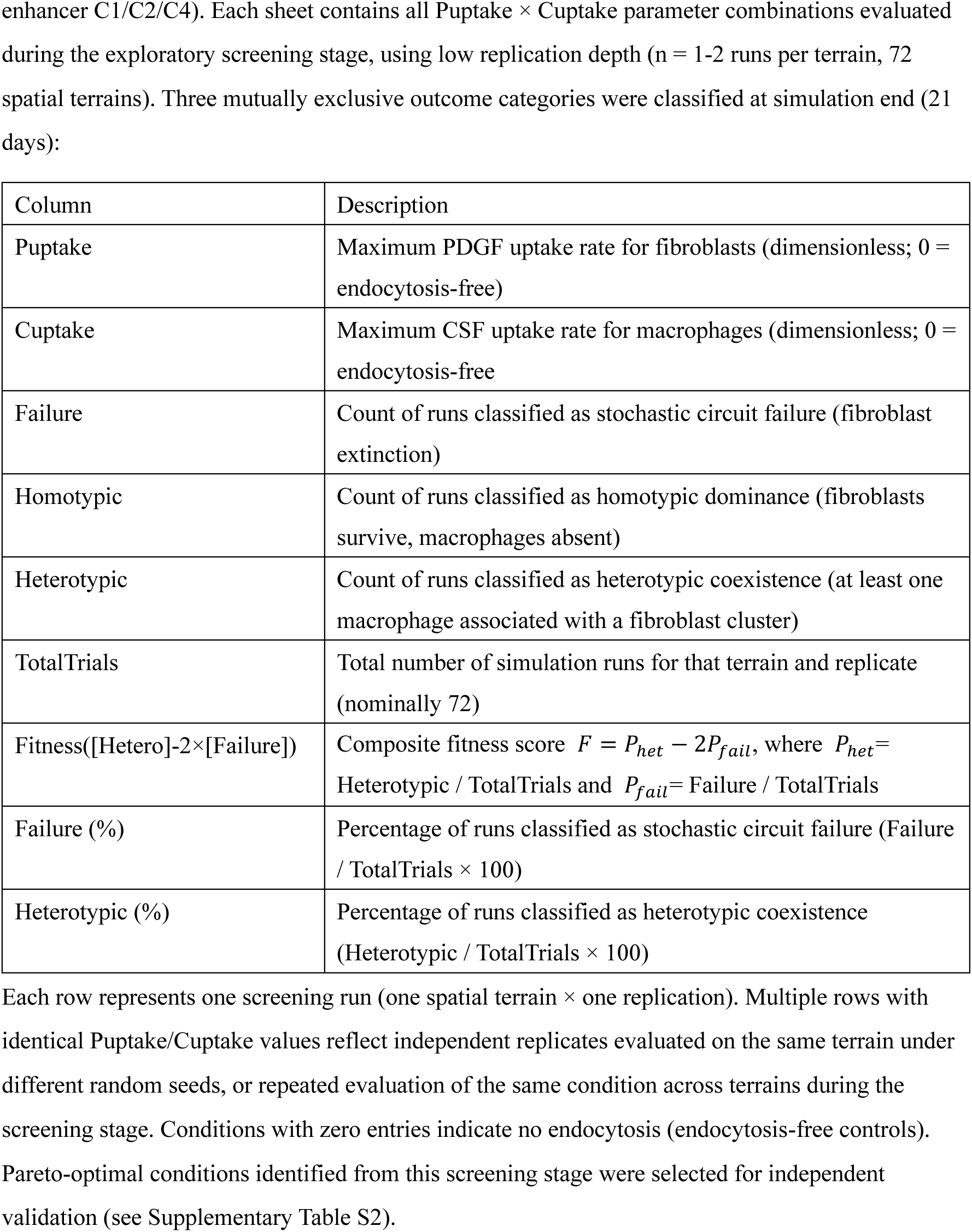
Pilot baseline and grid screening simulation results. Raw simulation outcome counts and summary statistics from the pilot baseline experiment (sheet: P1P1baseline) and grid screening stage (sheets: P1C1-P4C4) across nine enhancer conditions. Sheet: P1P1baseline Outcomes from the hypothetical homotypic baseline architecture in which both cell populations (CellA and CellB) produced and responded exclusively to PDGF, without CSF dependence. Simulations were performed under endocytosis-free conditions (PuptakeA = PuptakeB = 0) across 72 independent spatial terrains, with 12 stochastic replicates per terrain (n = 12 × 72 = 864 total runs). Three mutually exclusive outcome categories were classified at simulation end (21 days):

| Column | Description |
| --- | --- |
| PuptakeA | PuptakeA: PDGF uptake rate for CellA (dimensionless; set to 0 for all baseline runs) |
| PuptakeB | PuptakeB: PDGF uptake rate for CellA (dimensionless; set to 0 for all baseline runs) |
| A and B terminated | Count of runs in which both CellA and CellB populations became extinct simultaneously |
| A or B terminated | Count of runs in which either CellA or CellB became extinct |
| A and B survived (separate cluster) | Count of runs in which both populations survived but remained spatially segregated |
| A and B survived (mingled cluster) | Count of runs in which both populations survived and formed spatially intermingled clusters |
| TotalTrials | Total number of simulation runs for that terrain (nominally 72) |
| A and B (%) | Percentage of runs with simultaneous extinction of both populations ($\text{A and B terminated} / \text{TotalTrials} \times 100$) |
| A or B (%) \| | Percentage of runs with extinction of either population ($\text{A or B terminated} / \text{TotalTrials} \times 100$) |
| Puptake | Maximum PDGF uptake rate for fibroblasts (dimensionless; 0 = endocytosis-free) |
| Cuptake | Maximum CSF uptake rate for macrophages (dimensionless; 0 = endocytosis-free) |
| Failure | Count of runs classified as stochastic circuit failure (fibroblast extinction) |
| Homotypic | Count of runs classified as homotypic dominance (fibroblasts survive, macrophages absent) |
| Heterotypic | Count of runs classified as heterotypic coexistence (at least one macrophage associated with a fibroblast cluster) |
| TotalTrials | Total number of simulation runs for that terrain and replicate (nominally 72) |
| Fitness([Hetero]-2×[Failure]) | Composite fitness score $F = P_{het} - 2P_{fail}$, where $P_{het}$ = Heterotypic / TotalTrials and $P_{fail}$ = Failure / TotalTrials |
| Failure (%) | Percentage of runs classified as stochastic circuit failure (Failure / TotalTrials × 100) |
| Heterotypic (%) | Percentage of runs classified as heterotypic coexistence (Heterotypic / TotalTrials × 100) |

**Supplementary Table S2.** Validation simulation results. Raw simulation outcome counts and summary statistics from the independent validation stage across nine enhancer conditions (sheets: P1C1-P4C4). Screening-stage simulations are excluded. Each selected Puptake × Cuptake condition was evaluated using 72 independent spatial terrains and 8 independent stochastic replicates per terrain (576 simulations per condition). Sheets: P1C1, P1C2, P1C4, P2C1, P2C2, P2C4, P4C1, P4C2, P4C4

| Column | Description |
| --- | --- |
| Puptake | Maximum PDGF uptake rate for fibroblasts (dimensionless; 0 = endocytosis-free) |
| Cuptake | Maximum CSF uptake rate for macrophages (dimensionless; 0 = endocytosis-free) |
| Failure | Count of runs classified as stochastic circuit failure (fibroblast extinction) |
| Homotypic | Count of runs classified as homotypic dominance (fibroblasts survive, macrophages absent) |
| Heterotypic | Count of runs classified as heterotypic coexistence (at least one macrophage associated with a fibroblast cluster) |
| TotalTrials | Total number of simulation runs for that terrain and replicate (nominally 72) |
| Fitness([Hetero]-2×[Failure]) | Composite fitness score $F = P_{het} - 2P_{fail}$ (primary analysis; k = 2) |
| Failure (%) | Percentage of runs classified as stochastic circuit failure (Failure / TotalTrials × 100) |
| Heterotypic (%) | Percentage of runs classified as heterotypic coexistence (Heterotypic / TotalTrials × 100) |
| Fitness([Hetero]-[Failure]) | Composite fitness score $F = P_{het} - P_{fail}$ (sensitivity analysis; k = 1) |
| Fitness([Hetero]-4×[Failure]) | Composite fitness score $F = P_{het} - 4P_{fail}$ (sensitivity analysis; k = 4) |

Each row represents one spatial terrain (1 terrain × 8 independent stochastic replicates; TotalTrials = 72 per terrain under nominal conditions). Rows are grouped by Puptake × Cuptake condition. The endocytosis-free control condition (Puptake = 0, Cuptake = 0) is included in each sheet for within-enhancer-condition comparison. Statistical comparisons between endocytosis-enabled conditions and the matched endocytosis-free control were performed using one-tailed Mann-Whitney U tests with Bonferroni correction (k = 4 comparisons per enhancer condition); results are reported in the main text and figures.

Supplementary Movie S1

S1 Movie. Agent-based simulation of heterotypic coexistence under high PDGF / low CSF enhancer conditions.

Time-lapse simulation of the fibroblast-macrophage two-cell circuit over 21 days (30,240 min) in a 2,000 × 2,000 µm domain. Simulation was performed under enhancer condition P4C1 (macrophage PDGF secretion target = 4,000; fibroblast CSF secretion target = 1,000) without endocytosis (Puptake = Cuptake = 0), using random seed 13. Red circles, fibroblasts; grey circles, macrophages; yellow circles, capillaries. Background color indicates oxygen concentration (red, high; yellow, low). The simulation illustrates the nucleation phase (initial transient decline in cell number), followed by clonal expansion around capillaries, and convergence to a stable heterotypic coexistence state in which fibroblast and macrophage clusters co-localize. P4C1 represents a ceiling-effect condition in which failure probability is intrinsically low regardless of endocytic activity (see main text); this movie therefore shows a representative outcome of the predominant circuit state under this enhancer combination.

Supplementary File S1. PhysiCell configuration file (PhysiCell_settings.xml).

Complete PhysiCell (version 1.13.1) configuration file in XML format specifying all simulation parameters used in this study. The file defines the two-dimensional simulation domain (2,000 × 2,000 µm; BioFVM mesh spacing 20 µm), time step settings (diffusion 0.01 min, mechanics 0.1 min, phenotype update 6 min), and total simulation duration (30,240 min; 21 days). Microenvironment substrate parameters are specified for CSF, PDGF, and oxygen, including diffusion coefficients, decay rates, and boundary conditions. Cell type definitions are provided for fibroblasts, macrophages, and capillaries, including volume, motility, secretion, uptake, and cycle parameters. The representative random seed (seed 36) included in this file corresponds to one example terrain; independent simulations were performed across 72 random seeds as described in the Materials and Methods. This file is provided to facilitate full model reproducibility in conjunction with Supplementary File S2.

Supplementary File S2. PhysiCell cell hypothesis rules (detailed_rules.txt).

Plain-text file specifying the signal-response rules governing cell behavior in PhysiCell rule-based format. Rules are defined separately for macrophage, fibroblast, and capillary cell types. For macrophages: oxygen suppresses necrosis rate (Hill response; half-max 0.1, Hill power 12); CSF stimulates cycle entry (Hill response; half-max 2, Hill power 2); CSF suppresses PDGF secretion target via negative feedback (Hill response; half-max 2, Hill power 4); CSF stimulates CSF uptake (endocytosis; Hill response; half-max 2, Hill power 2; maximum uptake rate shown here represents one representative condition). For fibroblasts: oxygen suppresses necrosis rate (Hill response; half-max 0.1, Hill power 12); PDGF stimulates cycle entry (Hill response; half-max 2, Hill power 2); PDGF stimulates PDGF uptake (endocytosis; Hill response; half-max 2, Hill power 2; maximum uptake rate shown here represents one representative condition). The maximum uptake rates for CSF (Cuptake) and PDGF (Puptake) were systematically varied across parameter grids during screening and validation as described in the Materials and Methods; the functional form and Hill parameters were held constant across all conditions. No rules are assigned to capillary cells, which function as immobile oxygen-secreting agents. This file is provided to facilitate full model reproducibility in conjunction with Supplementary File S1.

Supplementary File S3. Python automation script for sequential random seed execution (random_run.py).

Python script developed for automated sequential execution of PhysiCell simulations across random seeds 1–72. The script reads simulation parameters from a JSON configuration file (settings_path.json) specifying the PhysiCell executable path and settings XML path, accepts user-defined seed range and output directory via standard input, and iteratively modifies the random seed and output folder path in the PhysiCell configuration XML before launching each simulation run. Output directories are organized by seed number (RundomSeed_1 through RundomSeed_72).

