## SupplementaryFileS for "Endocytosis suppresses stochastic collapse in fibroblast-macrophage circuits under shared resource competition": SupplymentaryFileS3.py

```
import xml.etree.ElementTree as ET
from pathlib import Path
import subprocess
import sys
import json

# 定数
RANDOM_SEED_XPATH = "./user_parameters/random_seed"
SAVE_FOLDER_XPATH = "./save/folder"
CELL_POSITIONS_XPATH = "./initial_conditions/cell_positions"
CONFIG_FILE_NAME = "settings_path.json"

# 関数

def modify_and_save_settings(target_xml_path, output_dir, seed_value):
    """XML設定を変更し、ターゲットパスに上書き保存する"""
    # ターゲットファイルを読み込む
    tree = ET.parse(target_xml_path)
    root = tree.getroot()

    # random_seed を変更
    seed_element = root.find(RANDOM_SEED_XPATH)
    seed_element.text = str(seed_value)

    # save/folder を変更
    folder_element = root.find(SAVE_FOLDER_XPATH)
    folder_element.text = str(output_dir.resolve()) 
    
    # initial_conditions/cell_positions の enabled を false に設定
    cell_pos_element = root.find(CELL_POSITIONS_XPATH)
    cell_pos_element.set("enabled", "false")

    # ターゲットファイルに上書き保存
    tree.write(target_xml_path, encoding="utf-8", xml_declaration=True)
    
    return


def run_simulation(exe_path):
    """PhysiCellシミュレーションを実行する"""
    cwd = exe_path.parent
    result = subprocess.run(
        [str(exe_path)], 
        cwd=str(cwd),
        check=True,
        shell=False,
    )
    return
 
def load_config(config_path_str):
    """JSON設定ファイルを読み込む"""
    config_path = Path(config_path_str)
    if not config_path.is_file():
        print(f"[FATAL] Config file not found: {config_path.resolve()}")
        sys.exit(1)
        
    with open(config_path, 'r', encoding='utf-8') as f:
        config_data = json.load(f)
        
    required_keys = ["PhysiCell_exe_path", "PhysiCell_settings_xml_path"]
    if not all(key in config_data for key in required_keys):
        print(f"[FATAL] Config file must contain: {', '.join(required_keys)}")
        sys.exit(1)

    config_data['PhysiCell_exe_path'] = Path(config_data['PhysiCell_exe_path'])
    config_data['PhysiCell_settings_xml_path'] = Path(config_data['PhysiCell_settings_xml_path'])

    return config_data


def get_save_path_input(prompt, default=None):
    """ユーザーからパスを受け取る"""
    while True:
        if default:
            prompt_full = f"{prompt} (default: {default}): "
        else:
            prompt_full = f"{prompt}: "
            
        value_str = input(prompt_full).strip()
        
        if not value_str and default:
            return default
        elif value_str:
            return value_str
        else:
            print("  [Error] Please enter a path.")


def main():
    """メイン処理"""
    # 標準入力から random seed を取得
    while True:
        seed_start = int(input("start random seed: "))
        seed_end = int(input("end random seed: "))
        if seed_start <= seed_end:
            break
        else:
            print("  [Error] Start seed must not be greater than end seed. Please try again.")

    print(f"Seed range: {seed_start} to {seed_end}")
    
    # 設定ファイルの読み込み
    config_file_path = CONFIG_FILE_NAME
    config = load_config(config_file_path)
    PhysiCell_exe_path = config['PhysiCell_exe_path']
    PhysiCell_settings_xml_path = config['PhysiCell_settings_xml_path'] 

    # 設定値のチェック
    if not PhysiCell_exe_path.exists():
        print(f"[FATAL] PhysiCell executable not found: {PhysiCell_exe_path}")
        sys.exit(1)
    if not PhysiCell_settings_xml_path.exists():
        print(f"[FATAL] Target settings XML not found: {PhysiCell_settings_xml_path}")
        sys.exit(1)

    # 結果保存用親ディレクトリを作成
    results_dir_str = get_save_path_input("Enter the parent directory path for saving results: ")
    results_parent_dir = Path(results_dir_str)
    results_parent_dir.mkdir(parents=True, exist_ok=True)
    print(f"outputs directory: {results_parent_dir.resolve()}")

    # シードを変更しながら実行
    for seed in range(seed_start, seed_end + 1):
        print(f"\n--- Processing Seed: {seed} ---")
        
        run_dir = results_parent_dir / f"RundomSeed_{seed}"
        run_dir.mkdir(exist_ok=True)
        print(f"  Output directory: {run_dir.resolve()}")

        # 設定ファイルを上書き変更
        modify_and_save_settings(PhysiCell_settings_xml_path ,run_dir, seed)

        # シミュレーションを実行
        success_run = run_simulation(PhysiCell_exe_path)
        if not success_run:
                print(f"  [WARNING] Simulation for seed {seed} failed.")

    print("\n--- All runs finished ---")

if __name__ == "__main__":
    main()
```
